# Integrative multi-omics analysis reveals conserved hierarchical mechanisms of FOXO3 pioneer-factor activity

**DOI:** 10.1101/2021.04.06.438676

**Authors:** Abigail K. Brown, Sun Y. Maybury-Lewis, Ashley E. Webb

## Abstract

FOXO transcription factors are critical for cellular homeostasis and have been implicated in longevity in several species. Yet how these factors directly affect aging, particularly in humans, is not well understood. Here, we take an integrated multi-omics approach to identify the chromatin-level mechanisms by which FOXO3 coordinates transcriptional programs. We find that FOXO3 functions as a pioneer factor in human cells, directly altering chromatin accessibility to regulate gene expression. Unexpectedly, FOXO3’s pioneer activity at many sites is achieved through a two-step process, in which chromatin accessibility is initially reduced, then transitions to an open conformation. The direct FOXO3 network comprises chromatin remodelers, including the SWI/SNF remodeling complex, which we find is functionally required for FOXO3 activity. We also identify a novel secondary network of activator protein-1 (AP-1) transcription factors deployed by FOXO3, which orchestrate a neuronal-specific subnetwork. Together, this hierarchical FOXO3 pioneer network regulates key cellular processes including metabolism, proteostasis, epigenetics and proliferation, which must be tightly controlled under changing conditions that accompany aging.

## Introduction

Aging is characterized by a gradual functional decline across organ systems due to both environmental and genetic factors. The insulin/IGF/FOXO signaling pathway is evolutionarily conserved and has been linked to healthy aging across taxa. This pathway’s influence on longevity has been demonstrated in model organisms, including *C. elegans*, *Drosophila*, and mice^1–3^, in which mutations that increase FOXO activity result in lifespan extension^4^. In humans, single nucleotide polymorphisms (SNPs) found in FOXOs, and FOXO3 in particular, have been correlated with longevity in several independent populations^5–7^. FOXO transcription factors function downstream of the insulin/IGF receptor, whereby ligand binding activates downstream effectors including AKT, which phosphorylates and inhibits FOXO transcriptional activity^8,9^. In contrast, in low nutrient or low growth factor conditions or in the presence of stress, FOXOs translocate to the nucleus where they regulate a myriad of cellular processes that maintain or restore cellular homeostasis^9,10^.

The Forkhead family of transcription factors are defined by a conserved winged helix DNA binding domain. This family includes the FOXO transcription factors, which directly bind and regulate a well-characterized consensus sequence (GTAAACA) that is conserved across species^11^. Through their direct networks, FOXO transcription factors function as key homeostatic regulators in response to stress or low nutrients^9,12^. Despite many studies linking FOXO factors to longevity in model systems, how these factors function at the chromatin level to directly impact aging in humans is poorly understood.

Tight regulation of cellular identity, homeostasis, and healthy aging relies on coordinated interactions at the epigenetic level between transcription factors and chromatin remodelers across the genome^13–15^. Together, these regulatory factors orchestrate stable gene expression programs that retain the ability to respond to intrinsic or extrinsic cues. A subset of transcription factors, termed “pioneer factors”, have the unique ability to access compacted chromatin, increase accessibility in that region and nucleate the binding of other transcriptional regulators^16–18^. Pioneer factor activity is critical for initiating chromatin accessibility and gene regulation during development and cellular reprogramming^19,20^. The first transcription factor shown to have this ability was FOXA, a member of the forkhead transcription factor family. *In vitro* studies showed that FOXA could bind DNA in compacted nucleosome arrays and render it accessible for DNase I digestion. The winged helix structure of the forkhead DNA binding domain resembles the nucleosome binding domain of the H1 and H5 linker histones and confers nucleosome binding ability to these transcription factors^21^. Intriguingly, work on FOXO1 has shown that, similar to FOXA, it functions as an ATP-independent chromatin remodeler *in vitro*, and this function is attributable to regions outside of the DNA binding domain^13^.

Studies in *C. elegans, Drosophila,* and mice have been integral in defining the gene expression programs through which FOXO/DAF-16 promotes longevity and the stress response^2,11,22,23^. While *in vitro* studies show intrinsic chromatin remodeling activity by forkhead transcription factors, *in vivo* studies in nematodes show that the switch/sucrose non-fermentable (SWI/SNF) chromatin remodeling complex is required for gene regulation by FOXO/DAF-16^24^. This ATP-dependent chromatin remodeler regulates the compaction and decompaction of genetic material without disrupting its capacity for replication, repair and transcription^25^. The SWI/SNF family (also known as BAF in mammals) specializes in increasing DNA accessibility through nucleosome sliding or ejection and does so through coupling with specific transcription factors.

In this study, we performed a multi-omics time-course analyses of FOXO3 activity in human cells. We demonstrate that FOXO3 functions as a pioneer factor *in vivo* to directly alter the chromatin landscape at specific sites. Unexpectedly, we find that accessibility at many targets is initially decreased, followed by opening of the chromatin. Mechanistically, FOXO3 promotes activation of chromatin remodelers, and we show that the SWI/SNF remodeling complex is functionally required for FOXO3’s activity in human cells. This activity dynamically coordinates a gene expression program to simultaneously regulate specific signaling pathways, chromatin organization and the cell cycle. Further, we find that FOXO3 deploys AP-1 transcription factors to regulate a secondary network critical for neuronal function. Together, our findings elucidate for the first time how FOXO3 directly promotes longevity and stress-associated gene networks through changes in chromatin accessibility, and indirectly through its position at the apex of a transcription factor hierarchy.

## Results

### FOXO3 functions as a pioneer factor in human cells

The extent to which FOXO3 acts as a pioneer factor to increase chromatin accessibility *in vivo* remains an unanswered question. To address this question and determine the chromatin-level effects of FOXO3 activation, we used a doxycycline-controlled induction system to rapidly and specifically activate FOXO3. We used ATAC-seq (Assay for Transposase-Accessible Chromatin)^26^ to map the global chromatin accessibility changes that occur in human U87 cells after FOXO3 activation for 8 and 16 hours (Fig. 1a). Consistent with FOXO3’s known role in restraining the cell cycle and promoting a quiescent state^23,27–31^, induction of FOXO3 causes cell cycle exit by 16 hours (Fig. 1b-c). Thus, the 8 and 16 hour time points correspond to chromatin alterations that occur before the reduction of proliferation and in the final arrested state, respectively. Cell cycle exit was accompanied by morphological changes consistent with arrest (Extended Data Fig. 1a). The ATAC-seq datasets separated by principal component analysis (PCA) based on whether there was FOXO3 activation or not, indicating changes in chromatin accessibility after 8 hours of induction, preceding cell cycle exit (Fig. 1d). We interrogated the regions that were altered in response to FOXO3 activation and identified 55,140 differentially accessible (DA) sites after 8 hours of FOXO3 activation. We observed both increases and decreases in accessibility (25.93% opening and 74.08% closing, respectively) (Fig. 1e, f Supplementary table 1), indicating a rapid epigenomic response to activation of FOXO3. After 16 hours of FOXO3 activation we observed 47,222 DA sites, with a majority of these sites (83.83%) displaying increased accessibility with FOXO3 activation (Supplementary table 2). Interestingly, we observed there were a number of regions previously closing with FOXO3 activation at 8 hours becoming open after 16 hours of FOXO3 activation (256 gene regions Fig. 1f). Analysis of the genomic distribution of the DA sites showed that the chromatin changes occurred mainly in distal intergenic or enhancer regions (40.57% and 43.0% of sites at 8 and 16 hours, respectively) and intronic regions of the genome (Fig. 1g).

**Fig. 1.**
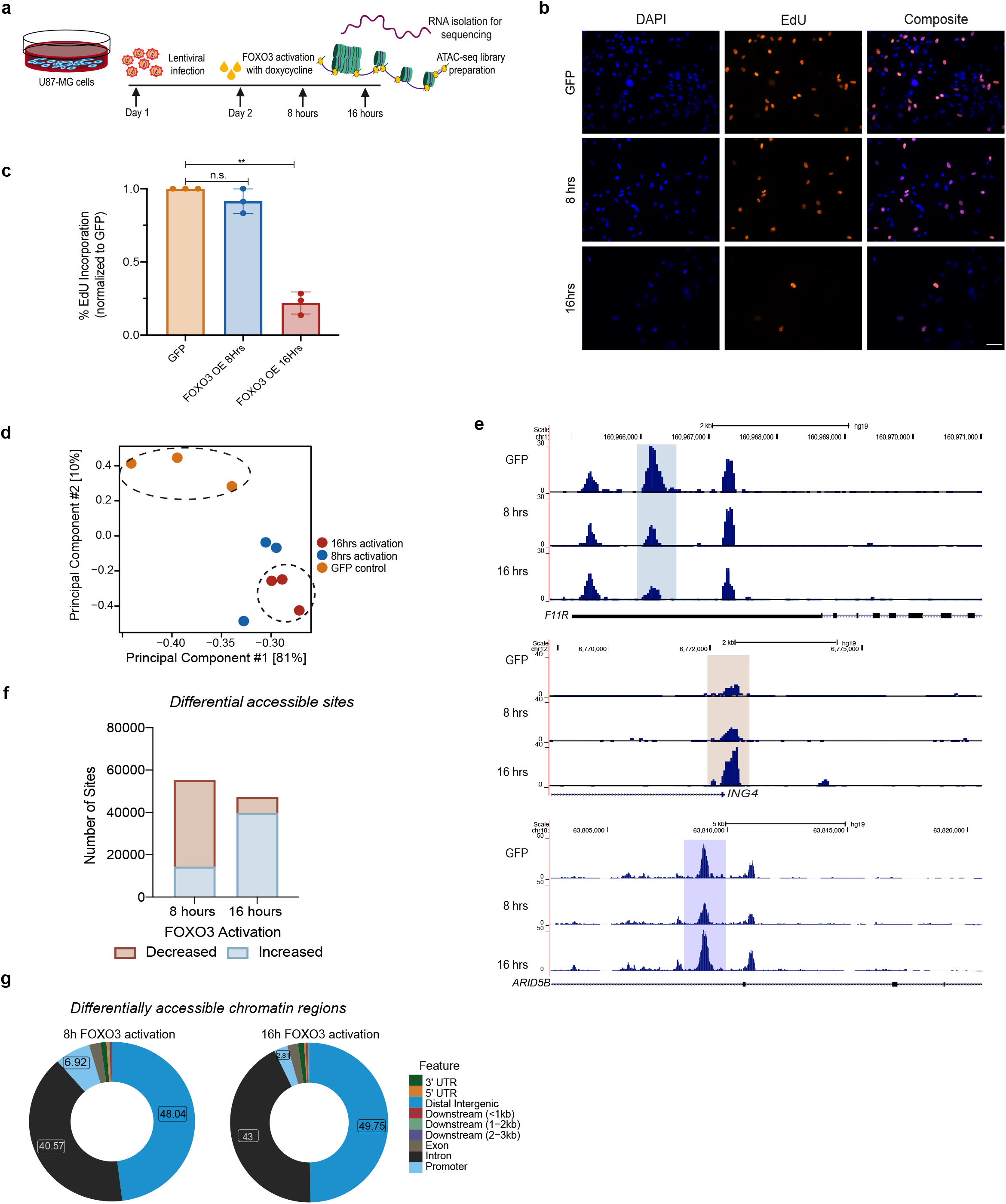
FOXO3 functions as a pioneer factor in human cells. **a)** Schematic of FOXO3 activation time course strategy used to define the transcriptome and chromatin accessibility changes induced by FOXO3. **b)** Representative images from EdU incorporation assays after FOXO3 activation for 0, 8, or 16 hours. Equal cell numbers were plated and EdU incorporation was performed for 2 hours. Scale bar, 160μm. **c**) Quantification of EdU positive cells described in (b) (*n*=3 biological replicates, \*\**p*<0.005, Student’s t-test). Error bars represent mean with standard deviation (SD). **d)** Principal component analysis (PCA) of ATAC-seq libraries after FOXO3 activation for 8 or 16 hours and GFP control. **e)** Distribution of the differentially accessible sites (increased or decreased accessibility) with FOXO3 activation at 8 hours and 16 hours. **f)** UCSC genome browser shots of example gene regions with altered chromatin accessibility over time with FOXO3 activation. Upper: decreased accessibility over time, middle: increased accessibility, bottom: decreased followed by increased accessibility. **g)** Donut graph shows the genomic distribution of the differentially accessible sites in 8 hours (*left*) and 16 hours (right) FOXO3-activated conditions. Promoter regions were defined as −1/+1kb around the TSS. Peaks were annotated based on distance from designated promoter region.

To determine the gene expression changes that accompany FOXO3-mediated chromatin remodeling, we performed RNA-seq using the same time course of FOXO3 activation (Fig. 1a). FOXO3 activation induced progressive transcriptional reprogramming, indicated by separation by principal component analysis (PCA) (Extended Fig.1b). When FOXO3 was activated for 8 hours we identified 2056 and 1408 genes that were significantly upregulated and downregulated, respectively (fold change ≥ 0.7 and a p value < 0.05). Using the same parameters, we observed similar numbers of genes after 16 hours of FOXO3 activation, with 2118 and 1340 genes upregulated and downregulated, respectively (Extended Fig. 1c, Supplementary Table 3). We used REACTOME pathway analysis to characterize the differentially expressed (DE) genes and observed significant enrichment for genes involved in FOXO3-mediated transcription, chromatin remodeling, and neuronal functions after 8 hours of induction, as well as the immune response and cell cycle after 16 hours of induction (Fig. 2a)^20^. Altogether, these analyses reveal that FOXO3 can function as a pioneer factor in human cells to rapidly induce transcriptional reprogramming.

**Fig. 2.**
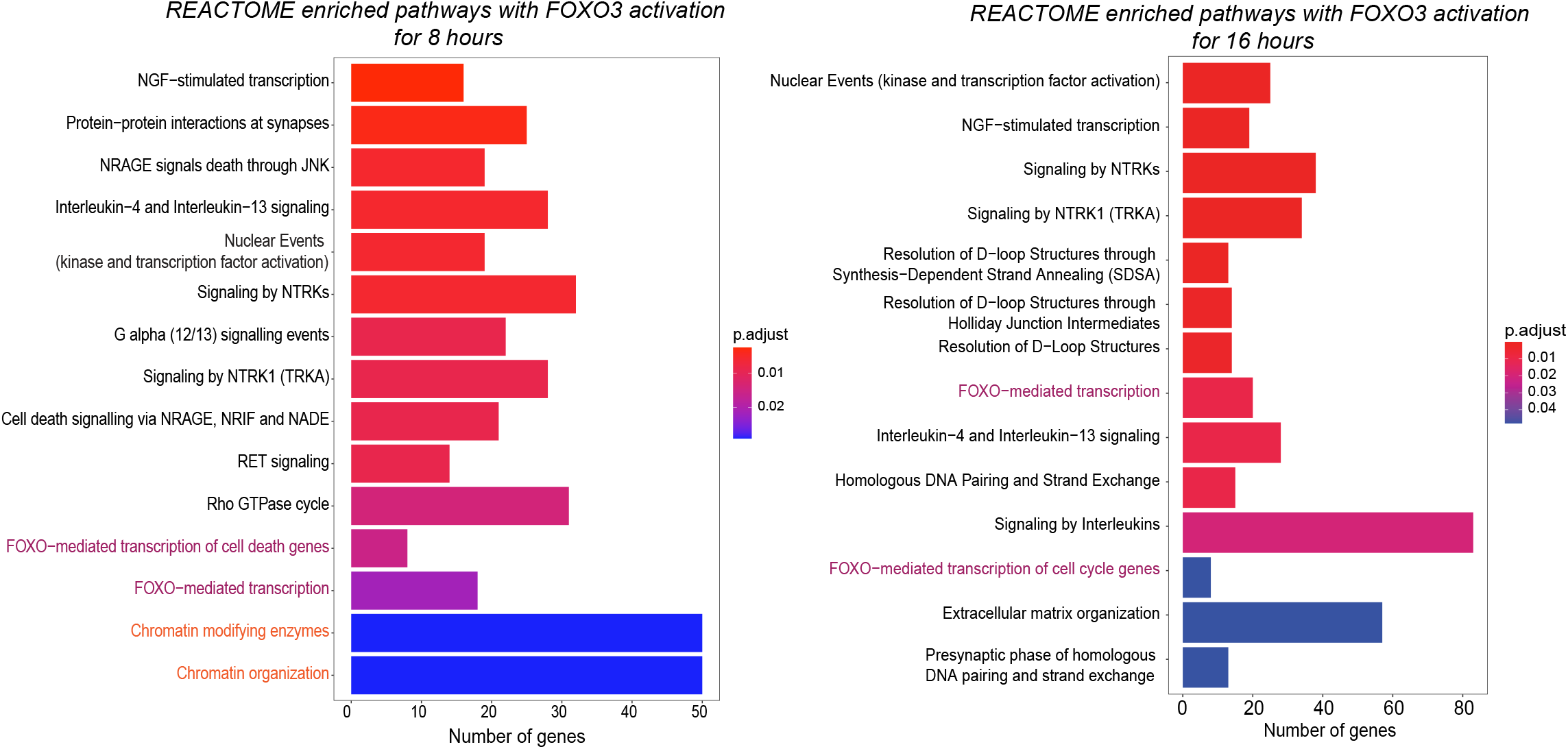
FOXO3 activation rapidly reprograms the transcriptome. REACTOME pathways analysis of differentially expressed genes between the GFP control and FOXO3 activation for 8 hours (*left*) and 16 hours (*right*).

### Identification of a direct FOXO3 target network in human cells that is conserved across evolution

To investigate the extent to which the chromatin accessibility changes observed in the ATAC-seq were a direct consequence of FOXO3 binding, we performed chromatin immunoprecipitation combined with deep-sequencing (ChIP-seq). Since suitable antibodies for FOXO3 in human cells are not available, we used the CRISPR epitope tagging ChIP-seq (CETCh-seq)^32^ method to integrate a FLAG tag into the endogenous FOXO3 locus in human U87 cells. We established two independent cell lines using this method (Fig. 3a) and validated each line by western blotting for the FLAG tag as well as the endogenous FOXO3 protein (Fig. 3b). Successful tag integration was further confirmed using immunocytochemistry for epitope-tagged and endogenous FOXO3, which showed co-localization and nuclear entry in response to PI3K inhibition (Fig. 3c). Thus, the established CETCh-seq cell lines express FLAG-tagged FOXO3 at endogenous levels, and epitope-tagged FOXO3 retains normal functionality in response to signaling cues.

**Fig. 3.**
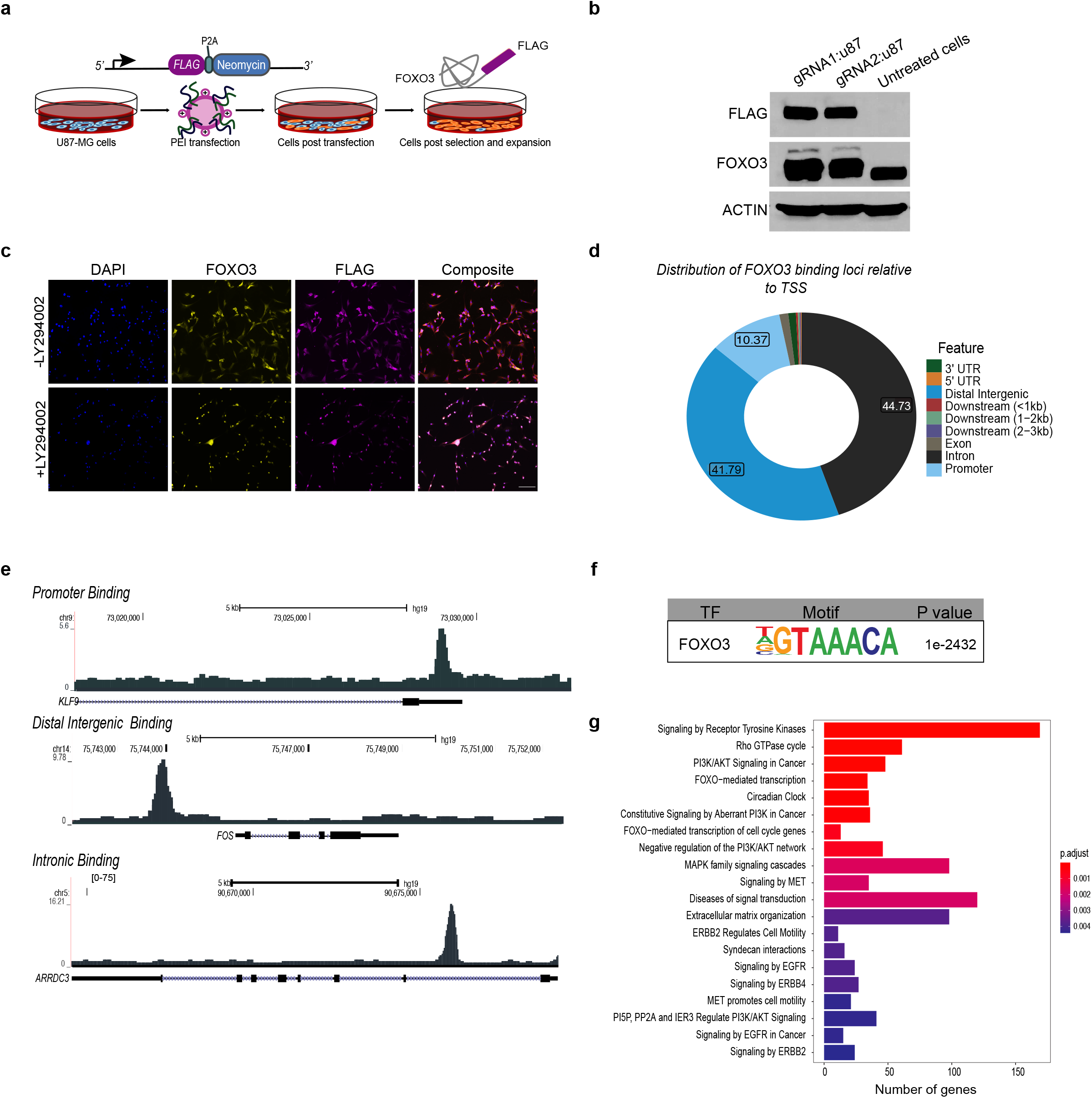
FOXO3 CETCh-seq identifies direct target genes in human cells. **a)** Schematic of the CETCh-seq experimental approach to establish a human cell line expressing epitope-tagged FOXO3 (FOXO3-FLAG) from the endogenous locus. **b)** Western blot showing validation of FLAG-tagged FOXO3 in two replicate cell lines. **c**) Representative images of U87 cell lines stably expressing FLAG-FOXO3 stained for endogenous FOXO3 (yellow) or FOXO3-FLAG (magenta). Cells are shown under basal conditions (−LY294002) or PI3K-inhibited conditions (+LY294002). Note the localization of the FOXO3 to the nucleus upon PI3K inhibition. Scale bar, 100μm. **d)** Donut graph shows the genomic distribution of all the FOXO3 binding sites. Promoter regions were defined as −1/+1kb around the TSS. Peaks were annotated based on distance from designated promoter region. **e)** FOXO3 binding at specified gene loci demonstrating binding at the TSS (*top*), distal intergenic (*middle*), and intronic (*bottom*) regions. **f)** Homer motif enrichment analysis of the called peaks following CETCh-seq. **g)** Bar plot shows significantly enriched REACTOME pathways (p. adjust < 0.05, Benjamin-Hochberg corrected) of the target genes of FOXO3. Bars are color coded according to the FDR corrected p value based on Benjamin-Hochberg analysis.

We next performed CETCh-seq on the established cell lines to identify the direct target genes of FOXO3. We used MACS2 to call significant peaks. The genomic distribution of FOXO3-bound sites was similar to previous studies in the mouse^11,23^, with the greatest enrichment for binding in intronic and enhancer regions, followed by binding directly at promoters (Fig. 3d-e). Motif analysis of the regions bound by FOXO3 shows significant enrichment for the forkhead box consensus sequence as the top binding motif (Fig. 3f), further validating the quality of the datasets. We used GREAT^33^ to assign the peaks to genes and found 2602 genes that are direct target genes of FOXO3 in human cells (Supplementary table 3). Gene ontology analysis revealed enrichment for expected functions such as FOXO3-mediated transcription and PI3K/AKT signaling (Fig.3g). Interestingly, we also observed novel pathways that are not typically associated with FOXO3 activity, such as Circadian Clock, Extracellular Matrix Organization, and Cell Motility. Together, these analyses identify the direct target network of FOXO3 in human cells and reveal strong enrichment for genes involved in regulating cellular and organismal homeostasis, including previously unknown functions.

FOXOs are conserved regulators of longevity and our previous work established that the conservation is at the level of direct targets^11^. To determine whether the evolutionary conservation extends to human, we performed a cross-species analysis comparing the FOXO3 target genes in humans with those previously found to be shared between multiple cell types in mice^11^. We observed a significant overlap between the direct target genes found in humans and mice, indicating strong evolutionary conservation of the network (Extended Data Fig. 2a, Supplementary Table 4; *p* = 7.02 × 10^−41^, Fisher’s exact test). REACTOME pathway analysis of these conserved genes revealed enrichment for pathways involving FOXO-mediated transcription, cell cycle regulation and chromatin modification. Interestingly, we also observed species-specific functions such as the cellular response to metals in human cells (Extended Data Fig. 2b-c). To determine whether human FOXO3 targets are also conserved in *C. elegans*, we performed a similar cross-species analysis with the DAF-16 targets previously identified^11,24^. We found 138 DAF-16/FOXO conserved targets genes between humans and *C. elegans* (Extended Data Fig. 2d, Supplementary table 5). These conserved target genes included well established targets of DAF-16/FOXO in *C. elegans* and are integral in regulating organismal homeostasis and longevity, such as *SOD-2, RICTOR, and FAT4 (sod-2, rict-1, and hmr-1* C. elegans orthologs respectively)^34–38^. Although the overlap did not reach statistical significance genome-wide, a number of REACTOME pathways were enriched among the shared genes, including the immune response and transcriptional regulation (Extended Data Fig. 2e). Collectively, these data indicate that a number of critical cis-regulatory targets of FOXO3 have been conserved from nematodes to humans.

To investigate the relationship between chromatin accessibility changes, direct binding of FOXO3, and regulation of gene expression, we performed an integrative analysis of the CETCh-seq, ATAC-seq and RNA-seq datasets. We first interrogated the DA sites for FOXO3 occupancy by CETCh-seq. We observed a significant overlap between FOXO3 binding and DA genes at 8 and 16 hours, indicating direct modulation of chromatin accessibility by FOXO3 (Fig. 4a). Specifically, FOXO3 binding was significantly enriched at 18.14% and 17.87% of DA sites at 8 and 16 hours, respectively. Further analysis of these FOXO3-directed DA genes showed a significant overlap between genes that are differentially accessible at 8 hours and 16 hours (Extended Data Fig. 3a). Integration of the RNA-seq data showed that while a number of genes induced by FOXO3 activation were direct targets (340, 31.89%), many DE genes were not occupied by FOXO3 (1976, 87.1%), suggesting induction of a secondary, indirect transcriptional network (Extended Data Fig. 3b). Interestingly, the direct program of induced genes was highly enriched for DA sites, indicating pioneer activity at these genes (65.88% of direct target genes had DA). In contrast, the secondary network of genes had fewer chromatin accessibility changes, with only 41.34% of DE genes associated with differential accessibility (Fig. 4b). Interrogation of the differentially expressed genes that were direct targets of FOXO3 revealed that FOXO3 directly regulated the expression of genes involved in transcription, response to metals and metabolic (leptin) signaling (Extended Data Fig. 3c). Thus, this multi-omics approach reveals for the first time a direct association between FOXO3 binding, altered chromatin accessibility, and transcriptional regulation, indicating chromatin-level pioneer activity in human cells.

**Fig 4.**
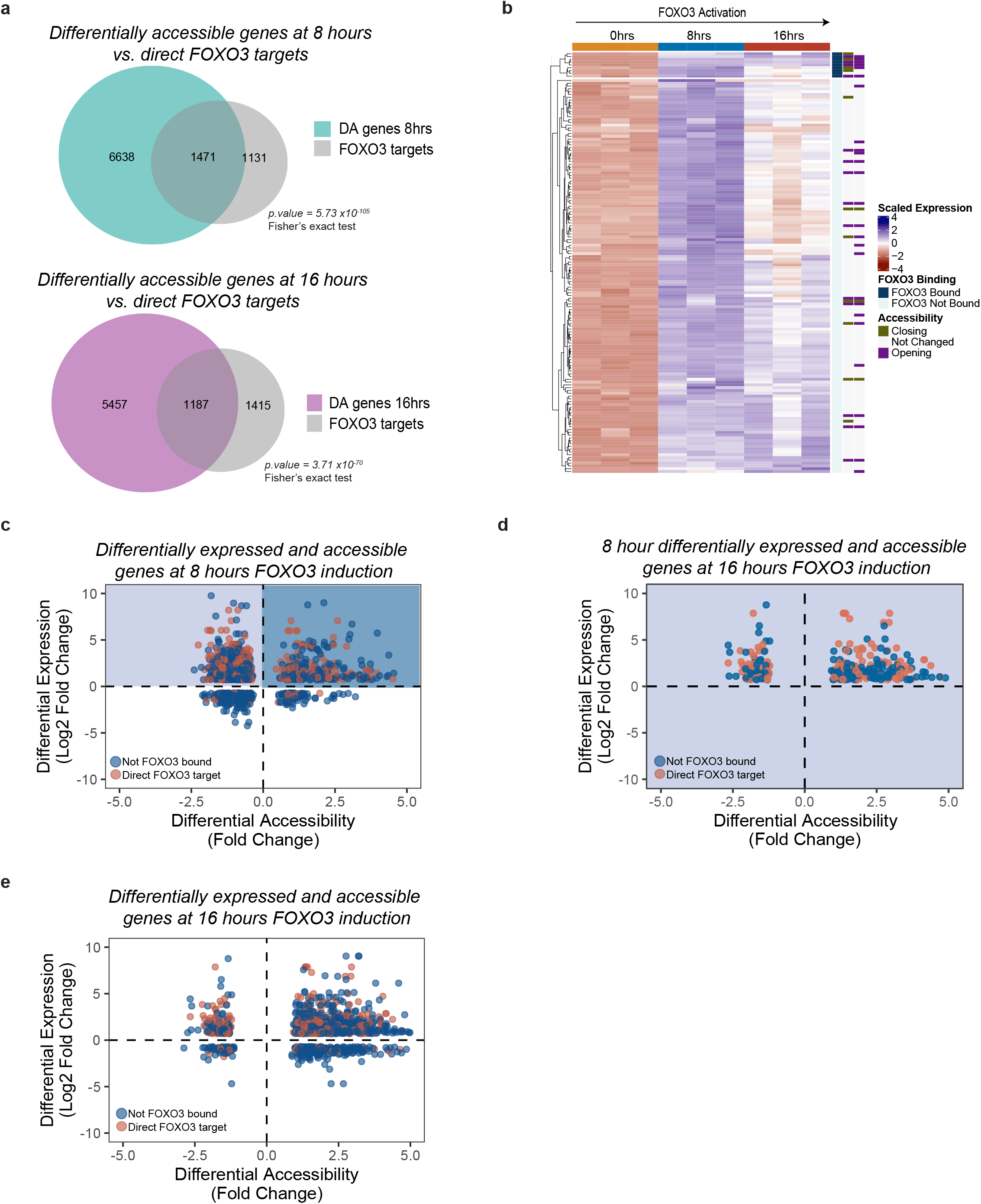
Integrated multi-omics analysis reveals two-step FOXO3 pioneer activity. **a)** Venn diagram depicting the overlap of genes directly bound by FOXO3 and genes at differentially accessible sites after 8 (*top*) and 16 (*bottom*) hours of FOXO3 activation. Overlap between the FOXO3 direct targets and differentially accessible genes is statistically significant at 8 hours (*p =* 5.73 × 10^−105^, Fisher’s exact test) and 16 hours (*p =* 3.71 × 10^−70^, Fisher’s exact test) of FOXO3 activation. **b)** Heatmap of the top 100 differentially expressed genes between GFP control and FOXO3 activated conditions. Rows show expression z score, samples are in columns. Annotation bars indicate direct target genes of FOXO3 as well as the chromatin accessibility status. **c)** Scatterplot showing fold change in chromatin accessibility versus fold change associated with gene expression (FDR < 0.05). Each dot represents a differential accessible site that is associated with a differentially expressed gene, with genes that are direct targets of FOXO3 colored in orange and others are colored in blue. **d)** Sites that are associated with chromatin closing but upregulation at 8 hours (upper left quadrant in (c)) are remapped at 16 hours, indicating increased accessibility following initial chromatin closing. **e)** Scatterplot showing the fold change in chromatin accessibility and gene expression of the genes at 16 hours.

To gain further insight into the chromatin dynamics underlying differential gene expression, we investigated the correlation between transcriptome and accessibility changes. As expected, we found that 8 hours of FOXO3 activation resulted mostly in gene activation, and many of these genes displayed increased chromatin accessibility (Fig. 4c; upper right quadrant). Surprisingly however, we observed that induction of gene expression was often associated with decreased accessibility after 8 hours of FOXO3 induction, followed by chromatin opening and sustained upregulation at 16 hours (Fig. 4c-d; 1f). This occurred with genes that were direct targets of FOXO3 (orange) and those that were not (blue). In general, after 16 hours of activation, increased gene expression was mostly associated with increased chromatin accessibility (Fig. 4e). This suggests that transcriptional activation is accompanied by rapidly increased accessibility at some sites, but many induced genes undergo a two-step chromatin restructuring in which chromatin is initially decreased in accessibility, possibility due to occlusion by remodeling factors, followed by conformational accessibility.

### FOXO3 induces a chromatin remodeling program in human cells

Our observation that FOXO3 alters chromatin accessibility in human cells through distinct mechanisms prompted us to investigate the network of remodelers associated with this activity. Since chromatin modifying enzymes and chromatin organization factors were among the top categories of genes induced by FOXO3 (Fig. 2), we interrogated the FOXO3 network for chromatin remodeling factors. Interestingly, the timeline of chromatin opening between 8 and 16 hours corresponds to significant changes in the expression of chromatin remodelers (e.g. ING4 and ARID2) and histone modifying enzymes (e.g. lysine demethylases KDM3A and KDM7A) (Fig. 5a). A number of these genes were directly regulated by FOXO3 and their chromatin states are altered by FOXO3 activation. Among the chromatin remodelers, we found that members of the BAF (mammalian SWI/SNF) complex were directly regulated by FOXO3, both in terms of their expression and their chromatin states (Fig. 5a). SWI/SNF is a multi-subunit ATP-dependent remodeling complex that functions together with DNA-binding transcription factors at enhancer regions. The complex includes both core components and complex-specific subunits, which are expressed to varying degrees in U87 cells (Fig. 5b). We identified two complex-specific genes, *ARID1A* and *ARID2*, that are directly regulated by FOXO3. Intriguingly, FOXO3 activity has opposite effects on these subunits: it upregulates *ARID2*, while it represses *ARID1A* expression, suggesting a subunit switching mechanism directed by FOXO3 (Fig. 5a). To determine whether FOXO3 activity is dependent on SWI/SNF chromatin remodelers, we induced FOXO3 activity in the presence or absence of a specific SWI/SNF (BAF) complex pharmacological inhibitor, PFI-3^39^. Using PFI-3 to inhibit the ATPase domain of the complex, Brg/Brm (Fig. 5c), we measured cell cycle exit in response to FOXO3 activation. We observed that BAF inhibition did not affect the cell cycle at 0 our 8 hours of FOXO3 induction, but inhibited FOXO3-induced cell cycle exit after 16 hours of activation (Fig. 5d-e, Extended Data Fig. 4a-d). Thus, FOXO3 activity reprograms the transcriptional network of chromatin and epigenetic regulators, suggesting SWI/SNF subunit switching, and ATP-dependent chromatin remodeling by SWI/SNF is required for FOXO3’s functional activity in human cells.

**Fig. 5.**
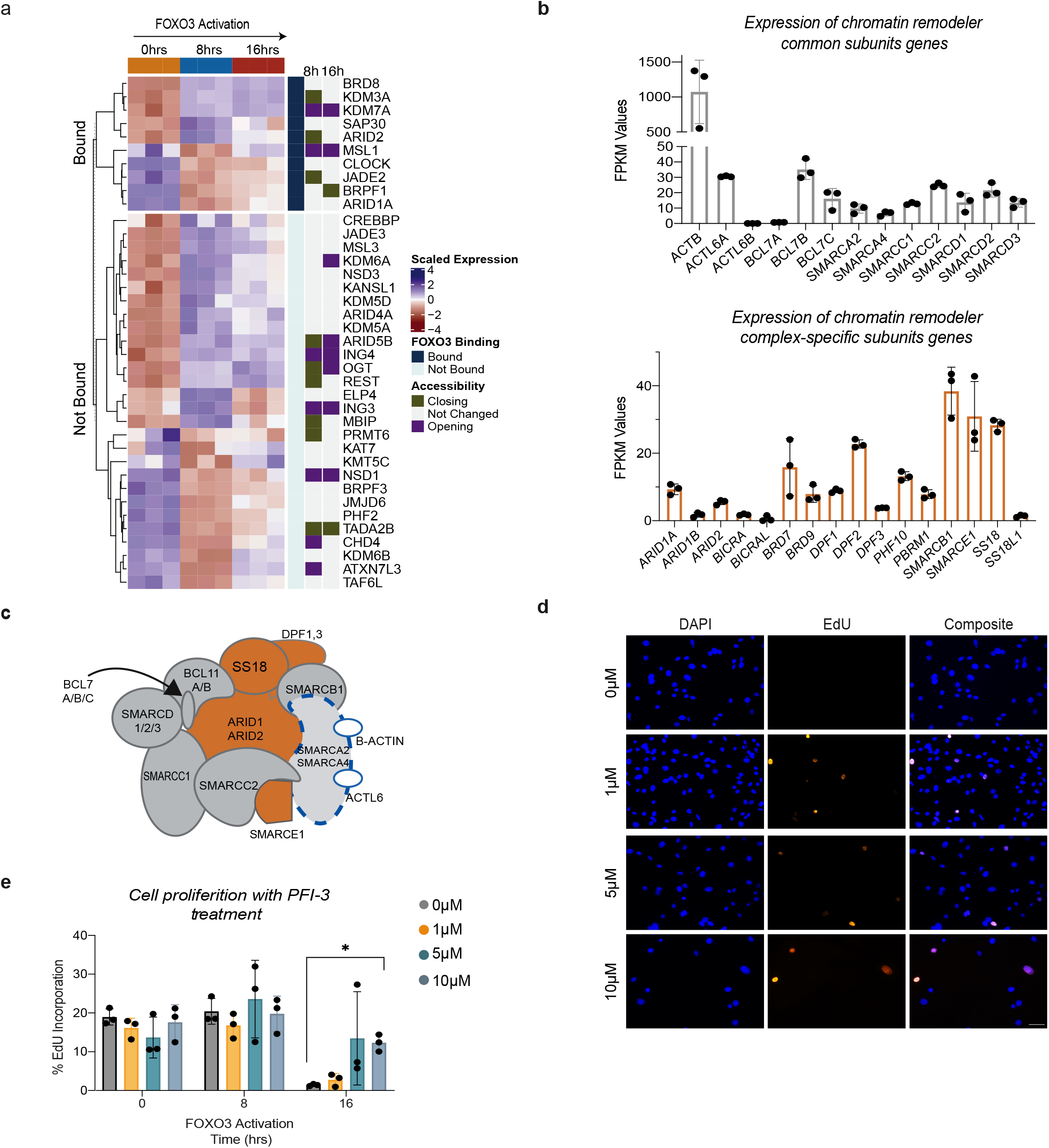
The BAF (SWI/SNF) chromatin remodeling complex functions with FOXO3 to promote cell cycle exit. **a)** Heatmap depicting the differential expression of genes enriched in the chromatin organization and chromatin modifying enzymes categories (from Fig. 2). Rows show expression z score, samples are in columns. Annotation bars show whether the genes are direct target genes of FOXO3 as well as the chromatin accessibility status. **b)** Transcript levels (FPKM values) of chromatin remodeler genes that comprise the mammalian SWI/SNF complex. Expression levels of the common subunits (*top*), complex specific members (*bottom*) are under basal conditions. Error bars represent mean with SD. **c)** Schematic of the mammalian BAF (SWI/SNF) complex. SMARCA2/SMARCA4 (Brg/Brm) is the catalytic subunit inhibited by PFI-3 and is highlighted by the dashed blue outline. **d)** Representative images of the EdU incorporation assay performed on FOXO3 activated cells treated with 5μM PFI-3 for 0, 8 and 16 hours (full set of images shown in Ext Fig. 4). Scale bar 160 μm. **e)** Quantification of the EdU incorporation assay in cells with FOXO3 induction and treatment with the BAF complex inhibitor, PFI-3 (*n* = 3 biological replicates, *p* = 0.0167, Student t-test). Error bars represent mean with SD.

### FOXO3 deploys an AP-1 pioneer sub-network

Our finding that many genes that are upregulated and changed in accessibility status upon FOXO3 activation are not direct FOXO3 targets raises the question of whether FOXO3 induces a secondary network of transcriptional regulators. To identify these putative transcription factors, we performed an *in silico* motif analysis of the DA sites that are not bound by FOXO3. Using the Homer differential motif discover algorithm^40^, we uncovered strong enrichment for binding motifs for the Activator Protein 1 (AP-1) family of transcription factors in DA sites after both 8 and 16 hours of FOXO3 activation (Fig. 6a). AP-1 transcription factors have been shown to regulate a wide range of cellular processes including growth, differentiation, and apoptosis. This family including the Fos (*FOS*, *FOSB*, *FOSL1*, *FOSL2*) and Jun (*JUN*, *JUNB*, *JUND*) protein families, as well as the closely related Activating Transcription Factors (*ATF2*, *ATF3*, and *BATF*) and Jun Dimerization Partners (*BATF3*/*JDP1* and *JDP2*). The proteins function as homo or heterodimers and bind DNA through their conserved bZIP domains^41–43^. We first investigated whether AP-1 family members are transcriptionally regulated by FOXO3. Analysis of the AP-1 family members revealed that *FOS*, *FOSL2*, *JUN*, *JUNB*, and *JUND* were all expressed in U87 cells and induced by FOXO3 at the RNA and/or protein levels (Fig. 6c-d). We also observed that both *FOS* and *JUNB* are direct targets of FOXO3, with FOS displaying increased chromatin accessibility with FOXO3 activation (Fig. 6b, e). Moreover, direct regulation of AP-1 factors by FOXOs appears to be evolutionarily conserved, as FOXOs also bind these factors in mouse tissues (Extended Data Fig. 5a)^11^. Functionally, the deployment of this secondary AP-1 network induced a neuronal-specific program, including synaptic functions, RET signaling and NGF-stimulated transcription (Fig. 6f). Interestingly, these functions are a subset of the FOXO3-directed transcriptional program (Fig. 2). Together these data indicate that FOXO3 induces a network of AP-1 transcription factors to drive a lineage-specific neuronal identity program upon cell cycle exit. Moreover, SWI/SNF chromatin remodeling activity is essential for the FOXO3-induced cell cycle arrest that restructures the chromatin landscape for the deployment of AP-1 factors (Fig. 6g).

**Fig. 6.**
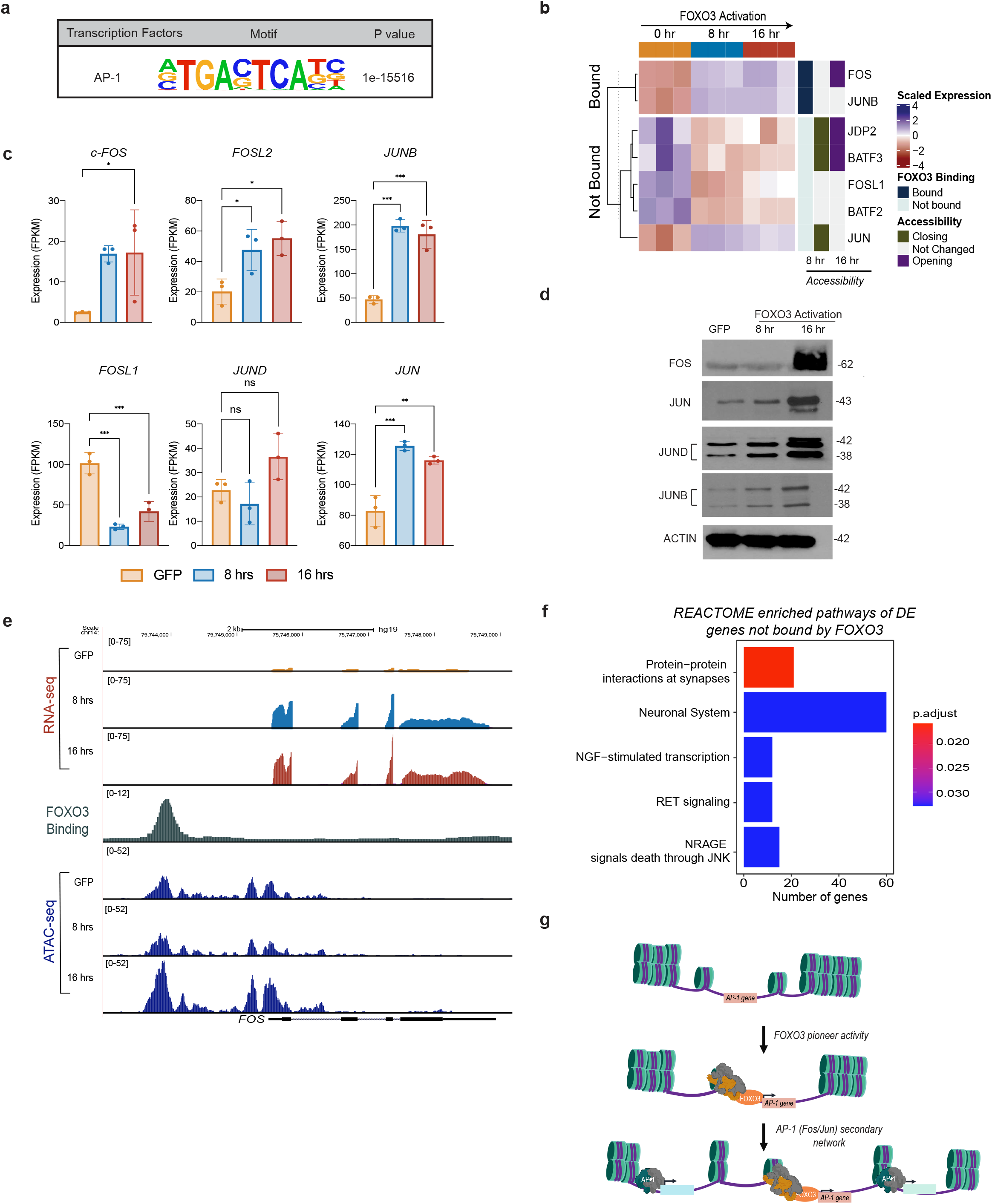
FOXO3 induces an AP-1 transcription factor network to drive a neuronal-specific transcriptional program. **a)** The AP-1 motif is highly over-represented at differentially accessible regions of the genome with FOXO3 activation at 8 and 16 hours. **b)** Heatmap of the differentially expressed AP-1 transcription factors in GFP control and FOXO3 activated samples at 8 and 16 hours. Rows represent expression z score, samples are in columns. Annotation bars indicate direct target genes of FOXO3 and chromatin accessibility status. **c)** Bar plots showing the transcript levels (FPKM values) of the AP-1 genes in basal conditions (GFP control), and FOXO3 activated 8 and 16 hour conditions (*n*=3 biological replicates, \*\*\**p*<0.0005, \*\**p*<0.005, \**p*<0.05 one-way ANOVA with Bonferroni corrected multiple comparison analysis). Error bars represent mean with SD. **d)** Western blot analysis of FOS, JUN, JUND, and JUNB protein levels with FOXO3 activation. One representative experiment of 3 independent replicates shown. **e)** UCSC tracks showing changes that occur in chromatin accessibility, and expression directed by FOXO3 at the *FOS* locus. **f)** REACTOME pathway analysis of differentially expressed genes that are not direct targets of FOXO3. **g)** Schematic of model showing FOXO3’s regulation of chromatin accessibility and transcriptional changes assisted by the AP-1 transcription factor network.

## Discussion

FOXO3 is an integral regulator of cellular and organismal homeostasis, with a conserved role in lifespan extension^4,9,11,44^. Seminal studies from the Cirillo and Zaret labs identified the Forkhead family of transcription factors, including FOXO1, as ATP-independent pioneer factors *in vitro*^13,21,45^. Work in *C. elegans* linked FOXO/DAF-16 activity with chromatin regulation, aging, and the stress response in invertebrates. Here, we used a multi-omics approach to define the chromatin-level mechanisms by which FOXO3 orchestrates a highly integrated transcriptional network to support processes that are critical for healthy aging, including epigenetic maintenance, cell cycle, neuronal function, metabolism, and cellular quality control.

While it has been established that FOXO3 is critical for cellular homeostasis, the mechanisms by which it dynamically regulates direct and indirect transcriptional programs to support key cellular functions are not well-defined. To address this gap in knowledge, we combined the recently developed CETCh-seq tool together with ATAC-seq and RNA-seq in human cells^32^. Our integrated analysis shows that FOXO3 directly alters the chromatin landscape as a pioneer factor to regulate a myriad of cellular processes including transcription/chromatin regulation, neuronal signaling pathways and the cell cycle^46–48^. We found that this direct pioneer network is evolutionarily conserved in mouse cells^11^ and numerous human FOXO3 genes were DAF-16 targets in *C. elegans*^11,22,24^. Targets that have been conserved from worms to humans include genes that regulate the immune response, TOR signaling, endosomal trafficking, oxidative stress, and transcription. We found similar results with the dFOXO targets from *D. melanogaster* data, with a smaller conservation of the target genes between human and files, including DUSP10, ARSJ, and RICTOR^49,50^. Thus, across evolutionary time, many of FOXO3’s key functions have been preserved.

FOXOs function as sensor transcription factors, responding to changing environments to allow cells to adapt to stress, nutrient availability, or aging. Our study reveals how FOXO3 activation rapidly alters the chromatin landscape to respond to molecular cues. We observed distinct mechanisms by which FOXO3 functions at the chromatin level. First, as described above, activation of FOXO3 led to transcriptional activation that could be linked to changes the chromatin landscape, with numerous regions of differential accessibility after 8 and 16 hours of activity. Second, and unexpectedly, we found that the majority of differentially accessible sites initially decrease in accessibility, despite being associated with upregulation of gene expression, followed by increased accessibility after 16 hours of FOXO3 activation. This finding suggests a two-step model of pioneer factor activity, whereby FOXO3 alters chromatin conformation and/or recruits remodeling factors that occlude accessibility to initiate gene expression, followed by establishment of a stable epigenetic state for maintenance of expression. Our analysis of chromatin remodelers showed that FOXO3 directly induces a number of genes that likely assist in the remodeling of chromatin state, including members of the SWI/SNF, MSL, and REST complexes. Further, inhibition of the SWI/SNF complex, suppressed the cell cycle exit phenotype previously observed with FOXO3 activation. Recent work has linked SWI/SNF (BAF) levels subunit exchange to cell cycle progression in development^51,52^, raising the possibility that FOXO3 may function by affecting SWI/SNF complex composition, as complex-specific subunits (ARIDs) were regulated by FOXO3 in our experiments.

Our study revealed that, while a significant portion of transcriptomic and epigenomic changes were directly controlled by FOXO3, there were numerous gene expression and accessibility changes that could not be accounted for by FOXO3 binding. Further analysis of these sites revealed enrichment for the activator protein 1 (AP-1) binding motif. AP-1 transcription factors are integral to cellular function and act through homo- or heterodimerization among family members^41,42^. The AP-1 factors are considered “immediate early genes” because they can be rapidly transcribed in response to various cellular stimuli. Our analysis revealed regulation of six AP-1 factors by FOXO3. Of this group, FOS was a particularly interesting target because it was induced at the transcript and protein levels, and its induction was accompanied by increased chromatin accessibility directly regulated by FOXO3. FOS and JUN family members were initially identified as oncogenes^53^, though they can also slow tumor growth, induce apoptosis, mediate neuronal activity, or initiate a nuclear stress response to promote recovery from neuronal injury^54–56^. AP-1 factors were also recently implicated in cellular senescence^57^ and work from invertebrate systems has linked FOXOs, AP-1 factors, and healthy aging, as Atf (CREB/*crh-1*) was identified as a DAF-16 regulated factor in worms^58,59^. Moreover, FOS has been shown to cooperate with dFOXO in *Drosophila* to regulate the DNA damage response^60^. Thus, the AP-1 factors are critical regulators of transcriptional programs in a number of cellular contexts. Our study reveals a novel hierarchical mechanism by which, concomitant with FOXO3-SWI/SNF-induced cell cycle exit, FOXO3 deploys the AP-1 factors to induce a secondary sub-network of gene expression that supports neuronal function in human cells.

In summary, our integrated analysis of chromatin and gene expression dynamics reveals the chromatin-level mechanisms by which FOXO3 functions as a pioneer factor to globally reprogram the human epigenome. This study provides novel mechanistic insight into FOXO3’s role in longevity, cellular homeostasis, and epigenetic stability in human cells.

## Materials and Methods

### Cell Culture

U87-MG glioblastoma cells (purchased from American Type Culture Collection; ATCC) were cultured in DMEM medium with 10% fetal bovine serum (FBS) and 1% penicillin-streptomycin-glutamine (PSQ) (Thermo Fisher Scientific) at 37°C and 5% CO_2_. 293T cells were used for lentivirus production and were also purchased from ATCC and cultured in the same conditions.

### Lentivirus production and infection

Lentiviruses (FOXO3, GFP, and rtTA) were generated in 293T cells with the helper plasmids RSV, VSV-G and PMDL. Viral supernatants were collected 24 hours after transfection with calcium phosphate and U87 cells were infected with a 1:1 ratio of viral supernatant to fresh media as described.

### EdU Incorporation Assays

Cells were incubated for 2 hours with 10 μM EdU, followed by fixation for 10 minutes with 4% paraformaldehyde. EdU incorporation was detected using the Click-iT EdU Alexa Fluro Imaging kit (Thermo Fisher Scientific) according to the manufacturer’s guidelines. Cells were imaged using Zeiss fluorescence microscope and EdU positive cells were counted manually.

### RNA-seq

Total RNA from was isolated from 3.5 × 10^6^ cells with the RNeasy mini kit (Qiagen). Samples were collected in triplicate for each time point. RNA quantification was performed using a nanodrop. PolyA libraries were generated using the NEB Next Ultra II library preparation kit and sequenced by Genewiz.

### RNA-seq data processing

Transcripts were processed using the previously outlined new Tuxedo protocol^61^. Briefly, Illumina adapters were removed from RNA-seq reads using Trimmomatic (version 0.36). Reads were then aligned to the human reference genome, hg19, using Hisat2 (version 2.1.0). Aligned reads were then assembled using StringTie (version 1.3.4d) and the expression levels all genes and transcripts were estimated^61–64^. Differential expression of genes was determined using DESeq2, with FDR < 0.05 and absolute log_2_foldchange ≥ 0.7 was considered statistically significant^65^. Overrepresentation of REACTOME pathways among differentially expressed genes was performed with ReactomePA R package^66^. Heatmaps were generated using the ComplexHeatmap package in R^67^.

### ATAC-seq

ATAC-seq libraries were prepared as previously described^68^. Briefly, 50,000 cells per time point were collected and washed in 1X PBS and centrifuged at 500g at 4°C for 5 minutes. Three independent replicates were performed. After collection, cells were lysed and the nuclei extracted by incubation in cell lysis buffer (10 mM Tris-HCl, pH 7.4, 10 mM NaCl, 3 mM MgCl_2_, 0.1% IGEPAL CA-630) and then immediately centrifuged at 500g at 4°C for 10 minutes. Transposition was then performed on whole nuclei by resuspending the nuclei in 50 μl of Transposition mix (22.5 μl of 1X TD Buffer and 2.5 μl of Tn5 enzyme) and then incubation at 37°C for 30 minutes. DNA was then isolated using the MinElute PCR purification kit (Qiagen). DNA was amplified using 9 PCR cycles, and libraries were then purified using the MinElute PCR purification kit. Library quality assessment was performed using the Bioanalyzer (Aligent). Paired-end sequencing (2×150 bp) was performed with Illumina HiSeq sequencing platform.

### ATAC-seq data processing

ATAC-seq raw reads were trimmed with TrimGalore! (version 0.40.0) and was aligned to hg19 reference genome using Bowtie2 (version 2.2.5)^69^. Duplicate reads were marked and removed using Picard (version 1.88, https://broadinstitute.github.io/picard/) and SAMtools (version 1.3.1)^70^, respectively. Peak calling was performed, after ATAC-seq specific quality control steps^71^, using MACS (version 2.1.1) with the FDR threshold 0.05^72^. Differential accessible sites were determined using DiffBind^73^.

### CETCh-seq construct generation and cloning

Cloning and generation of the CETCh-seq constructs was performed as previously described^32^. The 3xFLAG-P2A-Neomycin epitope tagging donor construct was obtained from Addgene (pFETCh-Donor, plasmid # 63934). Homology arms targeting the human FOXO3 transcription factor was ordered as synthetic dsDNA genomic blocks (IDT), PCR amplified and then DNA was isolated using the Qiagen PCR purification kit (Qiagen). Assembly of the final donor plasmid was performed by Gibson Assembly (New England BioLabs). sgRNA targeting near the 3’-end stop codon of FOXO3 were designed using CHOPCHOP (http://chopchop.cbu.uib.no) to minimize off target effects. The pSpCas9(BB)-2A-GFP vector (Addgene plasmid # 48138) was cut with BbsI (NEB) restriction enzyme and then ligated with the designed gRNA.

### CETCh-seq generation of FOXO3-FLAG mammalian cell line

The pFETCh-Donor plasmid with homology arms targeting the *FOXO3* locus, along with Neomycin resistance, as well as CRISPR/Cas9 plasmid with targeting gRNA (targeting the STOP codon) were cotransfected into U87-MG cells. 1.0 × 10^5^ cells were plated in 12 well plates and allowed to adhere overnight. pFETCh constructs and Cas9/gRNA plasmids were cotransfected for all CETCh-seq experiments. Cells were transfected with 2:1 (pFETCh-Donor: Cas9/gRNA) ratio of plasmids, with a total DNA concentration of 10μg, using Polyethylenimine (PEI). U87-MG cells were selected with 250 μg/ml G418 for approximately 4 weeks after which the remaining cells were pooled into one well for an additional two weeks of selection.

### Immunocytochemistry

For immunocytochemistry, 50,000 cells were plated onto glass coverslips placed in 24 well plates. Cells were fixed for 10 minutes with 4% paraformaldehyde and permeabilized with 0.4% Triton-X in PBS. After blocking with 5% normal donkey serum (NDS), cells were incubated with primary antibody (FLAG, dilution 1:200, Sigma-Aldrich F1804; FOXO3, dilution 1:200, CST 75D8) for 2 hours and then with Alexa Flour secondary antibody (anti-mouse, 1:500; anti-rabbit, 1:500, Jackson Laboratories) for 1 hour. Nuclei were stained with 4,6-diamidino-2-phenylindole (DAPI), at 1:5000 dilution for 10 minutes. Imaging was carried out using Zeiss fluorescence scope with 40X objective.

### Western Blotting

Cells were lysed in RIPA buffer (50 mM TrisHCl pH 8, 150 mM NaCl, 1% Triton X-100, 0.5% sodium deoxycholate, 0.1% SDS, 10 mM NaF, 1 mM sodium orthovanadate, 1mM EDTA) on ice for 10 minutes. Protein concentration was determined using the Qubit Protein Assay Kit (Thermo Fisher Scientific). Equal amounts of extracts were mixed with 3X Lamelli buffer, samples were then diluted with normalizing amounts of 1X Lamelli buffer and boiled at 95°C for 5 minutes. Samples were loaded unto 10% bis-acrylamide gel and electrophoresed for 2 hours at 130 V in 1X Running buffer. Proteins were transferred unto nitrocellulose membranes with Mini-PROTEAN transfer system (Bio-rad). Membranes were blocked in 5% BSA/Tris Buffered Saline (TBS) for 1 hour at room temperature, then incubated with primary antibodies (Anti-FLAG and Anti-FOXO3, dilution 1:1000, Anti-Actin 1:5000) 5% BSA/TBS overnight at 4°C. Membranes were washed with TBS-T ween (TBS-T) and incubated with 5% milk/TBS at 1:15,000 dilution. Detection was performed using Pierce ECL Western Blot Substrate (Thermo Fisher Scientific).

### Chromatin immunoprecipitation

CETCh-seq U87-MG cells were treated with a PI3K inhibitor LY294002 for 1 hour. A total of 2 × 10^7^ cells were fixed with 1% formaldehyde for 10 minutes, quenched with 0.125 M glycine for an additional 5 minutes. Cells were then lysed using SDS lysis buffer (50 mM Tris-HCl pH 7.4, 10 μM EDTA, 1% SDS, mini-Protease inhibitor tablet) on ice for 5 minutes. Samples were collected and then centrifuged at 3500g for 10 minutes at 4°C. Lysis buffer was removed, and the cell pellet was resuspended in RIPA buffer (1% IGEPAL CA-630, 1% sodium deoxycholate, 1% SDS, 1 mini-Protease inhibitor tablet). Sonication was performed using Diagenode Bioruptor for 10 minutes. Soluble chromatin was obtained by centrifugation at 14,000rpm for 15 minutes at 4°C. A fraction of this soluble chromatin was taken from each of the samples as the input control for the experiment. Immunoprecipitation of chromatin was performed overnight using 5 μg of anti-FLAG (Sigma-Aldrich F1804) coupled to mouse Dyna beads IgG (Invitrogen) at 4°C. The beads were then pelleted and washed one time with low salt, two times with high salt, three times with LiCl wash buffers (Millipore Sigma), and two times with TE buffer (10 mM Tris-HCl, 1 mM EDTA). Immunoprecipitated DNA was eluted from beads using elution buffer (4% SDS, 0.1 M NaHCO_3_) at 65°C for 2 hours. The beads were removed from the samples and the crosslinks were reversed overnight at 65°C. DNA isolation was performed using 25:24:1 phenol:chloroform:isoamyl alcohol (Sigma Aldrich) followed by ethanol precipitation of DNA for 1 hour at −20°C (Input DNA was isolated using the same procedure). DNA was resuspended in 100μl of molecular grade water (Thermo Fisher Scientific). Libraries were generated by Genewiz and 2×150 bp paired-end sequencing was performed using an Illumina HiSeq. An average of 90 million reads were obtained per library.

### ChIP-seq data processing

Raw sequencing data files were trimmed using Trimmomatic (version 0.36) and aligned to the hg19 reference genome using Hisat2 (version 2.1.0)^62,63^. Duplicate reads were marked with Picard (version 2.9.2) and removed with SAMtools (version 1.9)^70^. MACS2 (version 2.1.1) was used to call peaks with a q value cutoff of 0.05^72^. Peaks were assigned to genes using GREAT, with the parameters to −25kb/+10kb around the transcription start sites (TSS)^33^. Motif analysis was performed using Homer (version 4.7) findGenomeMotifs.pl tool.

### Antibodies

The antibodies that were used in this study are as follows:

| Antibodies | Company | Product # |
| --- | --- | --- |
| FOXO3 | CST | 75D8 |
| FLAG | Sigma Aldrich | F1804 |
| HA | Roche | 11583816001 |
| JUN | Santa Cruz Biotechnology | sc-398615 |
| FOS | CST | 4384 |
| JUND | CST | 5000 |
| JUNB | Santa Cruz Biotechnology | sc-8051 |

## Supplementary Information

Supplementary Table 1: Differential accessible sites 8 hours post FOXO3 activation.

Supplementary Table 2: Differential accessible sites 16 hours post FOXO3 activation.

Supplementary Table 3: Results of differential expression with FOXO3 activation at 8 and 16 hours.

Supplementary Table 4: FOXO3 CETCh-seq target sites in human U87-MG cells.

Supplementary Table 5: List of all conserved FOXO3 targets in humans and mice.

Supplementary Table 6: List of all conserved FOXO3 targets in humans and *C. elegans*.

## Acknowledgments

We would like to thank Dr. Richard Freiman (Brown University) for critical reading of the manuscript. We would also like to thank the members of the Webb laboratory for suggestions on experimental design, and Shawn Williams for advice on bioinformatics and critical reading of the manuscript. These studies were funded by P20 GM109035 (PI: D. Rand), U19 AG023122 (MPI: S. Cummings, S. Melov) to A.E.W. A.K.B has received support from the IMSD training program at Brown University and an NIH supplement to promote diversity.

## Author contributions

A.K.B. and A.E.W. conceived and designed the study and experiments. A.K.B conducted the experiments under the supervision of A.E.W. S.Y.M-L. performed the processing and analysis of ATAC-seq data. A.K.B analyzed the RNA-seq and CETCh-seq data. A.K.B. and A.E.W wrote the manuscript.

## Extended Data

**Extended Data Fig. 1 Cellular and transcriptome changes upon FOXO3 activation. a)** Representative immunocytochemistry images of HA-tagged FOXO3 in cells where FOXO3 was activated at 8 hours (*middle*) and 16 hours (*bottom*) with GFP control (*top*). Scale bar represents 50 microns. **b)** PCA analysis of FOXO3 activated RNA-seq samples, FOXO3 activation 8, 16 hours and GFP control. **c)** Volcano plot showing fold changes for differentially expressed genes between GFP control and FOXO3 activation 8 hours (*left*) and GFP control and FOXO3 activation 16 hours (*right*).

**Extended Data Fig. 2 FOXO target genes are conserved in humans and mice. a)** Venn diagram depicting the overlap of the FOXO3 target genes in humans and the shared FOXO target genes in mice (p = 7.02 × 10^−41^, Fisher’s exact test). **b-c)** REACTOME pathway analysis of the FOXO target genes **b)** conserved between humans and mice or **c)** mouse specific (top) and human specific (bottom). **d)** Venn diagram depicting the overlap of the FOXO3 target genes in humans and DAF-16 target genes in *C. elegans* (p = n.s, Fisher’s exact test). **e)** REACTOME pathway analysis of conserved human FOXO3/worm DAF-16 target genes.

**Extended Data Fig. 3 FOXO3 pioneers chromatin accessibility and transcriptional changes. a)** Overlap of the differentially accessible genes that are direct targets of FOXO3 at both timepoints of FOXO3 activation (*p* < 2.2 × 10^−16^ Fisher’s exact test). **b)** Venn diagrams depicting overlap between FOXO3 binding and differential expression at 8 and 16 hours (p =8.21 × 10^−38^ and p = 9.04 × 10^−63^, respectively, Fisher’s exact test) **c)** REACTOME pathway analysis of genes that are differentially accessible (8 hour) and directly regulated by FOXO3.

**Extended Data Fig. 4 Inhibition of SWI/SNF restores proliferation in cells with high FOXO3 activity.** Representative images of the EdU incorporation assay performed on FOXO3 activated cells treated with **a)** 0 μM, **b)** 1μM **c)** 5μM **d)** 10μM PFI-3 along with FOXO3 activation for 0, 8 and 16 hours. Scale bar 160 microns.

**Extended Data Fig. 5 AP-1 regulation by FOXO3 is conserved in mice. a)** UCSC genome browser shots of FOXO binding across cell types in the mouse at the *Junb* and *Fos* mouse gene loci.

