## Supplemental Data 1 for "Integrative multi-omics analysis reveals conserved hierarchical mechanisms of FOXO3 pioneer-factor activity": Extended Data Figures.pdf

**Extended Data Fig.1**

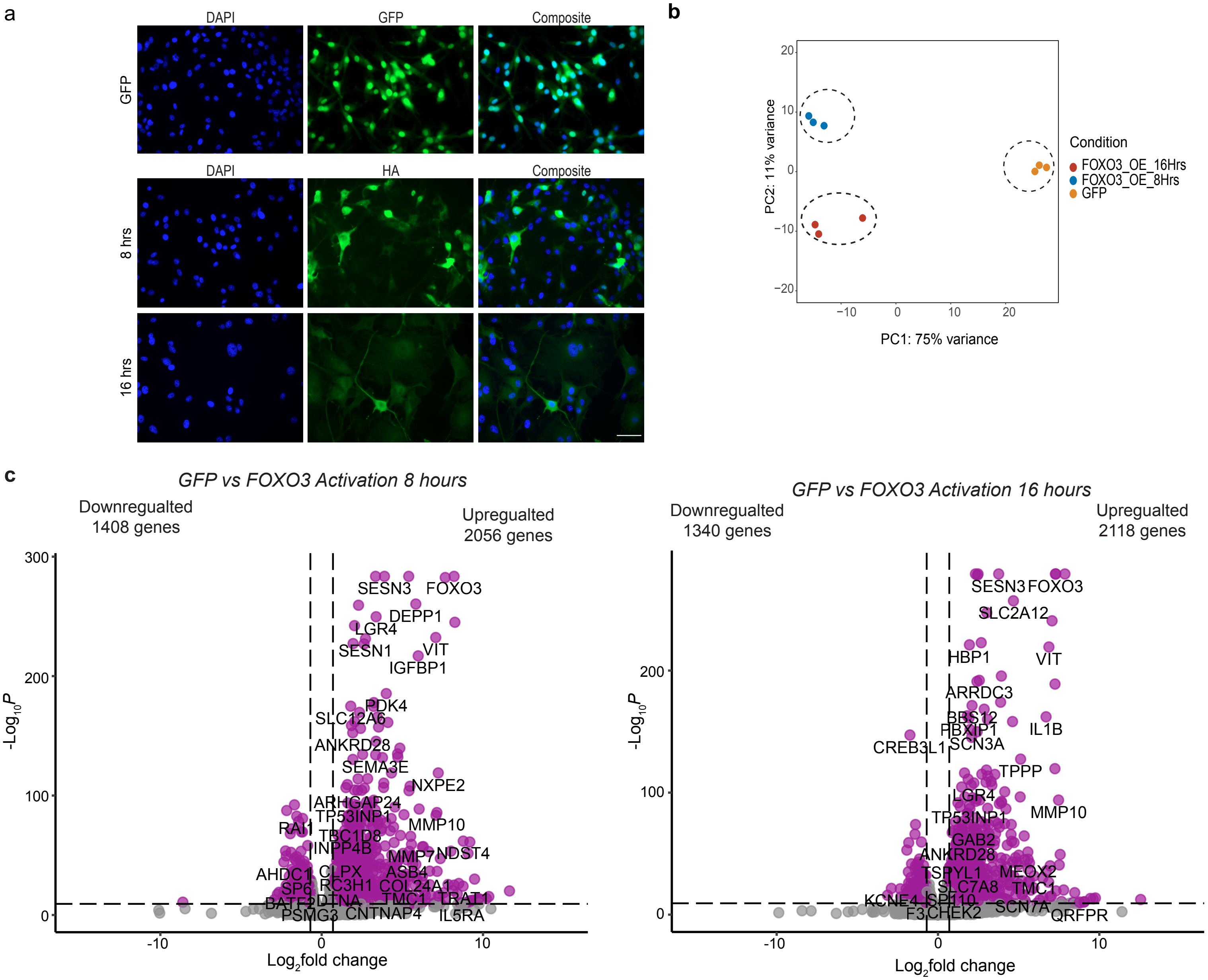

**Extended Data Fig. 2**

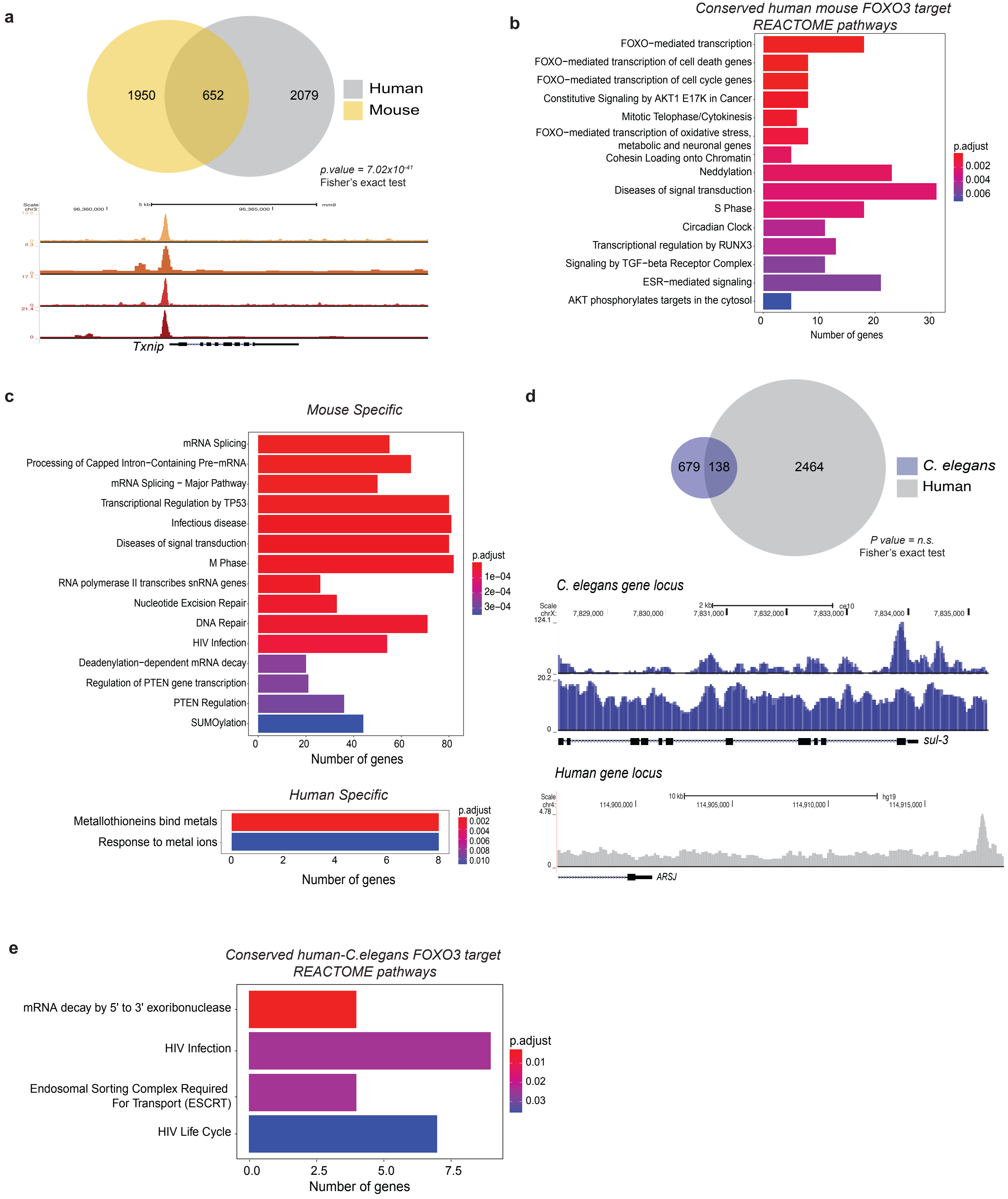

**Extended Data Fig.3**

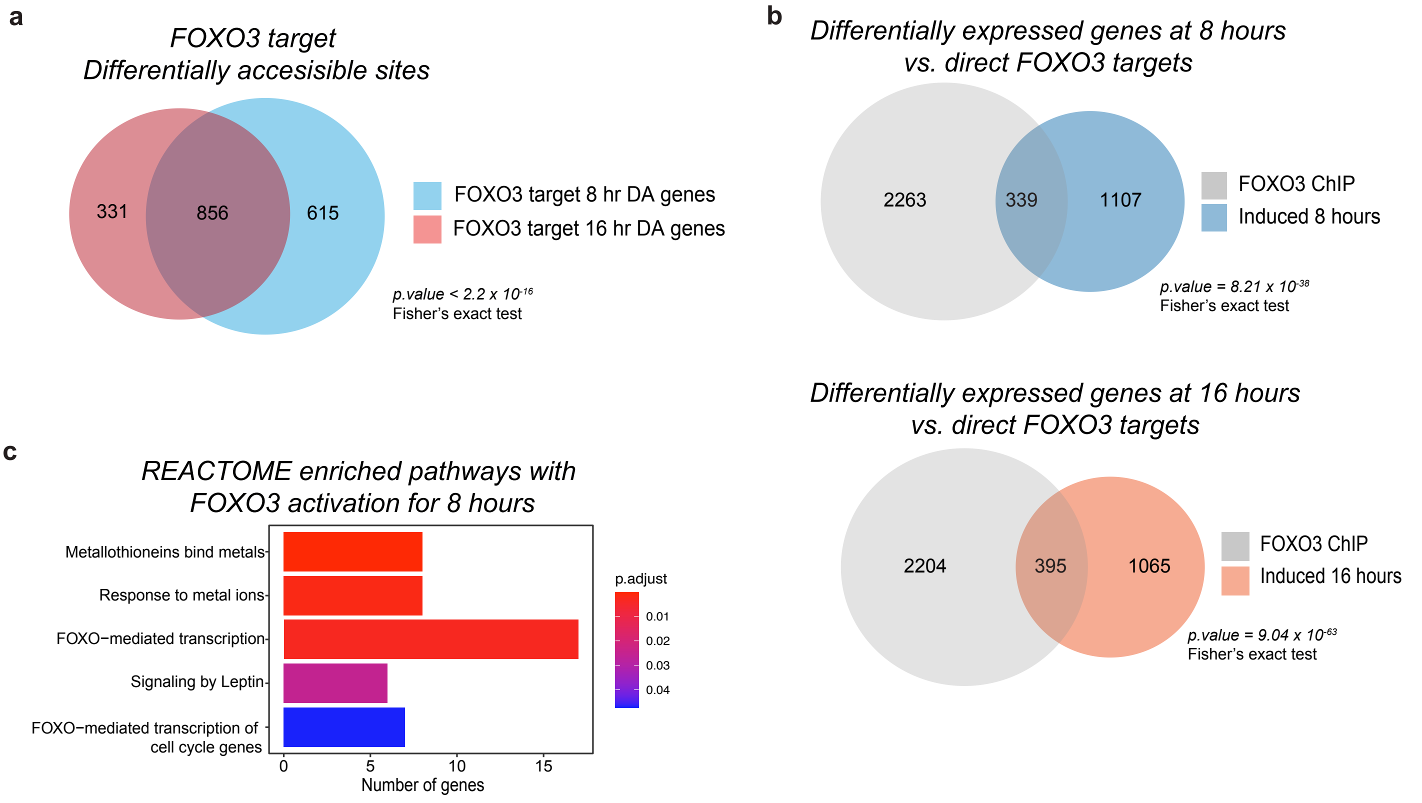

Extended Data Fig.4

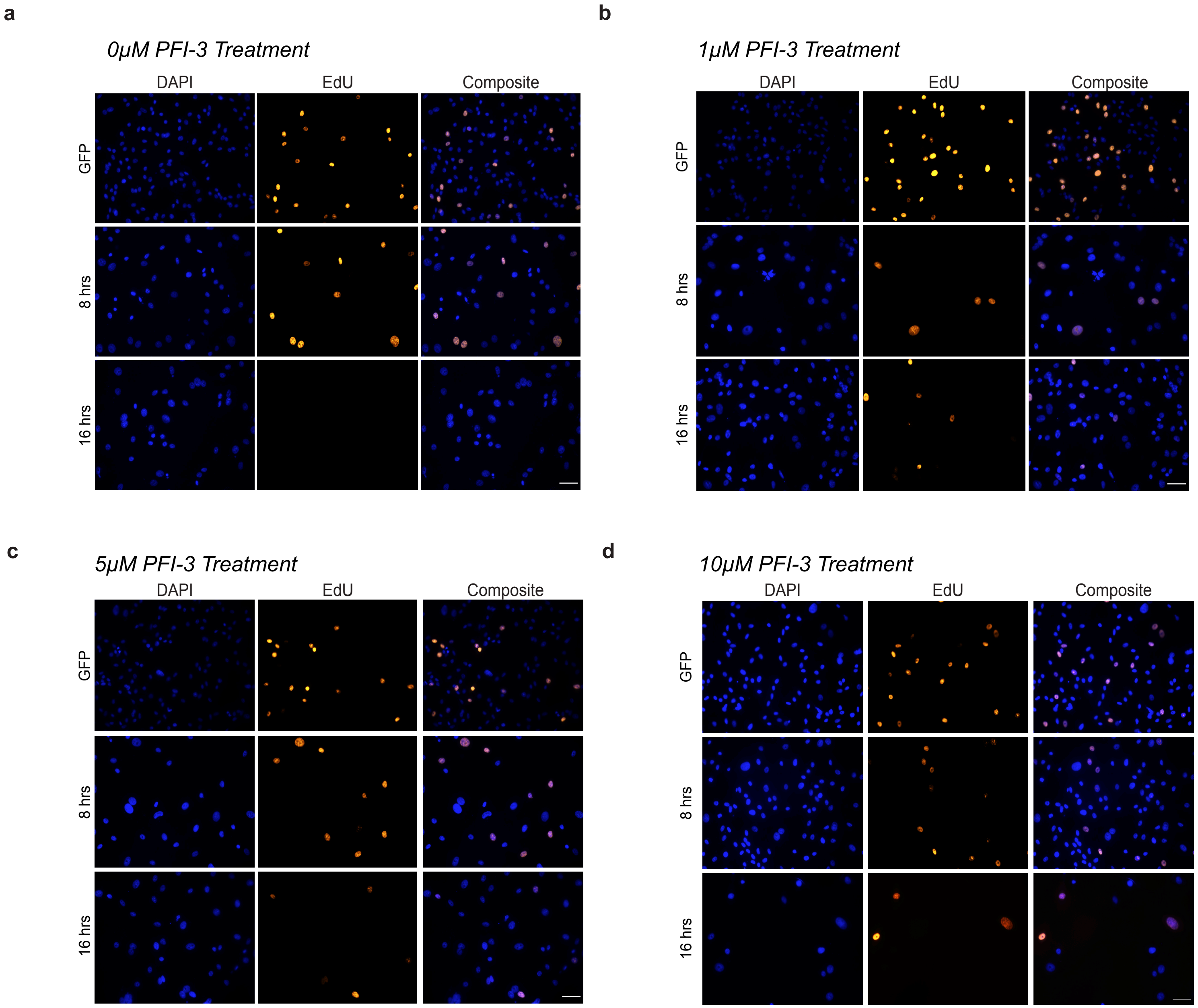

### Extended Data Fig. 5

**a**

Mouse *Junb* locus

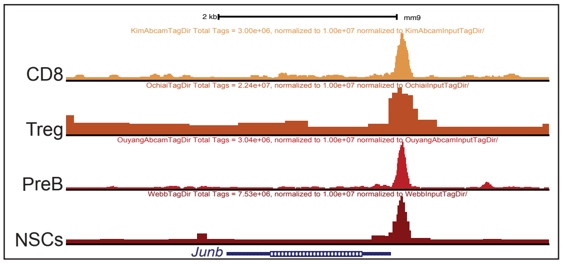


Mouse *Fos* locus

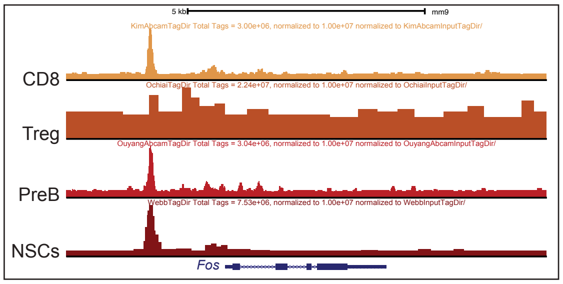
